# Trem-2 modulation of gut microbiota is blunted during hepatotoxic injury and uncoupled from liver repair responses

**DOI:** 10.1101/857078

**Authors:** Inês Coelho, Nádia Duarte, Maria Paula Macedo, Carlos Penha-Gonçalves

**Author notes:** Corresponding author: Carlos Penha-Gonçalves, Address: Instituto Gulbenkian de Ciência (IGC), Rua da Quinta Grande n°6, 2780-156 Oeiras, Portugal. **Authors contributions** Inês Coelho: study concept and design. Acquisition, analysis and interpretation of data, drafting of the manuscript; Nádia Duarte: acquisition, analysis and interpretation of data, drafting of the manuscript. Maria Paula Macedo, study concept and design, drafting of the manuscript; Carlos Penha-Gonçalves: study concept and design, drafting of the manuscript and study supervision. All authors approved final submission.

## Abstract

The involvement of gut microbiota in liver disease has been addressed in the context of the “leaky gut hypothesis” postulating that dysbiosis allow microbial components to elicit liver inflammatory responses and hepatic tissue damage. Conversely, commensal gut microbiota acting on innate immune receptors protect against hepatotoxic insults. Given that mice deficient for the triggering receptor expressed on myeloid cells-2 (Trem-2) show increased vulnerability to experimental drug-induced hepatic damage we explored the possibility that Trem-2 is a modulator of gut microbiota composition.

We found that microbiota composition in untreated Trem-2 KO mice differs from the wild-type showing overall decrease in microbiota diversity and increased representation of Verrucomicrobia. Interestingly, induction of liver damage with hepatotoxic drugs blunted this microbiota diversity difference and altered phyla composition with increased representation of Verrucomicrobia during acute hepatic injury and Proteobacteria during chronic challenge. Furthermore, co-housing experiments that homogenized microbiota diversity showed that the increased liver tissue vulnerability to hepatotoxic insults in Trem-2 KO mice was not dependent on microbiota composition. This work uncouples Trem-2 dependent alterations in gut commensal microbiota from Trem-2 pro-recovery effects in the damaged liver tissue. These findings support the possibility that unlinked actions of innate immune receptors contribute to disease association with microbiota alterations, particularly with the Verrucomicrobia phylum.

**Importance:** Trem-2 is a mammalian innate immunity receptor involved in development and resolution of tissue damage, namely in the brain and in the liver. Nevertheless, it is not known whether gut microbiota is contributing to these Trem-2 mediated phenotypes. We found that Trem-2 KO mice spontaneously display different gut microbiota composition as compared to wild-type mice, namely with increased abundance of the phylum Verrucomicrobia. Notably these differences do not impact the control of Trem-2 on liver tissue vulnerability to hepatotoxic insults. This work uncouples Trem-2 modulation of gut microbiota and the role of Trem-2 on responses to liver damage. This work brings new insights on role of innate immune receptors on the association of organic and systemic diseases with gut microbiota.

## Introduction

Susceptibility to inflammatory disease often depends on the interplay of host genetic factors with both immune system activation and microbiota composition^1^. Clinical evidence and experimental models have shown that disruption of gut homeostasis leads to leakage of bacterial products that reach the liver fueling pro-inflammatory responses^2^. Yet, it is still unknown how dysbiosis causes liver disease beyond increasing intestine permeability. The unexpected increased vulnerability of germ-free mice to hepatotoxic treatments^3^ raises the possibility that a “symbiotic” crosstalk of the microbiota with the innate immune system is critical in liver resilience against hepatocytic insults. Counter-intuitively, innate immunity sensor pathways (e.g. Myd88) have a beneficial effect in the response to toxic-induced liver damage^3^. This suggests that innate stimulation confers a tolerogenic environment that conditions the pro-inflammatory reactivity of the liver innate immune cells limiting tissue damage responses. A liver-protective role of the axis gut microbiota-liver innate cells is supported by early evidence showing that intestinal microbiota swiping leads to significant reduction in the number of liver mononuclear cells^4^. Furthermore, it has been shown that germ-free mice have delayed response to partial hepatectomy supporting the possibility that interaction of microbiota components with innate immunity receptors expressed in mononuclear cells play a role in liver regeneration^5^.

Liver macrophages may have either detrimental or beneficial effects in the response to liver damage and during tissue repair^6–9^. In steady-state conditions, the liver resident population of Kupffer cells predominantly exerts tolerogenic functions, namely by stimulating the suppressive activity of regulatory T cells. Such suppressor roles can be overcome through increased TLR signaling, highlighting the role of innate immunity receptor signaling in the modulation of liver responses to gut-derived microbial products^10^.

The triggering receptor expressed on myeloid cells (TREM) gene family, in particular the Trem-2 receptor, play decisive roles in tuning the activation threshold of macrophages and in directing phagocytic responses^11^. Initial studies showed that Trem-2 is expressed on newly differentiated macrophages and acts to restrain macrophage activation^12^. More recently, Trem-2 has been intensively studied in the context of neurodegenerative diseases with genetic variants identified as disease susceptibility factors in human populations^13^. In Alzheimer’s disease, expression of Trem-2 was suggested to control the phagocytic capacity and survival of microglia determining distinct cell activation states upon disease induction^14, 15^. Interestingly, these results were also recapitulated in models of multiple sclerosis and aging^15, 16^. In the liver, we recently found that expression of Trem-2 promotes macrophage restorative activities during recovery from acute and chronic liver damage^17^. Additionally, a recent study showed that antibiotic treatment followed by CCl4-induced acute liver injury, abolished the increase in hepatic enzymes levels observed in Trem-2 KO mice, while intestinal permeability was not affected^18^.

Here we investigated whether Trem-2 liver regeneration mechanisms are impacted by alterations in the gut microbiota. Analyzing Trem-2 KO mice we found that Trem-2 modulation of gut microbiota composition is not determining the outcomes of acute and chronic liver injury.

## Materials and Methods

### Mice and experimental models

All procedures involving laboratory mice were in accordance with national (Portaria 1005/92) and European regulations (European Directive 86/609/CEE) on animal experimentation and were approved by the Instituto Gulbenkian de Ciência Ethics Committee and the Direcção-Geral de Veterinária (the Official National Entity for regulation of laboratory animals usage). Trem2-deficient mice^19^ were kindly provided by Marco Colonna, Washington University School of Medicine, St. Louis, MO. C57BL/6 mice and Trem-2 KO mice were bred and housed under a 12-hr light/dark cycle in specific pathogen free housing facilities at the Instituto Gulbenkian de Ciência.

To induce acute liver injury, C57BL/6 and Trem-2 KO male mice with 10 weeks of age were fasted for 15 hours prior to intra-peritoneal injection with 300mg/Kg of acetaminophen (N-acetyl-p-aminophenol (APAP)) (Sigma, St. Louis, MO, USA) in PBS or PBS only. Liver was collected 3 days after injection.

In the model of chronic liver fibrosis and fibrosis regression, C57BL/6 and Trem-2 KO males with 7-8 weeks of age received PBS or 20%v/v carbon tetrachloride (CCl4, Sigma, St. Louis, MO, USA) in olive oil, administered at 0,4mL/Kg, twice a week during 4 weeks by intra-peritoneal injections^20^. Liver was collected at day at day 3 post-injection (fibrosis regression).

For co-housing experiments, wild-type and Trem-2 KO mice were co-housed at weaning and left together for 5 weeks. After 5 weeks of co-housing mice were either treated with APAP or separated for further 5 weeks.

### Histology

Histological analysis were performed in the Histopathology Unit of Instituto Gulbenkian de Ciência. Livers were fixed in 10% formalin and embedded in paraffin. Non-consecutive 3µm sections were stained with hematoxylin-eosin and examined under light microscope (Leica DM LB2, Leica Microsystems, Wetzlar, Germany). Necrosis was blindly assessed by a trained pathologist. Necrosis scoring: 0: no necrosis; 1: single cell centrilobular necrosis to centrilobular necrosis without central to central bridging necrosis; 2: centrilobular necrosis with central to central bridging; 3: centrilobular necrosis with central to central bridging and focal coalescent foci of necrosis; 4: centrilobular necrosis with central to central bridging with large coalescent foci of necrosis. Images were acquired on a Leica DM LB2.

Intestines were fixed by immersion in methacarn solution (60% methanol, 30% chloroform, 10% acetic acid) for 48 hr, followed by two successive washes each in methanol for 35 min, ethanol for 30 min, and xylene for 25 min. Tissue samples within cassettes were then submerged in melted paraffin at 68°C for 1 hr, removed, and kept at room temperature until sectioning. Paraffin blocks were cut into 4-mm-thick sections, deparaffinized and stained with hematoxylin-eosin.

### Fecal sample collection and DNA extraction

Fresh stool samples were collected at each indicated time point and immediately frozen and stored at −80°C. DNA was extracted from stool samples using a combination of the QIAamp DNA Stool Mini Kit (QIAGEN) following the manufacturer’s instructions and a mechanical disruption, where bacterial membrane disruption was enhanced using 0.1mm glass beads in ASF buffer (QIAGEN) and high-speed shaking in a TissueLyzer device (2 minutes, 30Hz; QIAGEN). Samples were stored at −20°C.

### Library preparation and sequencing

Bacterial DNA extraction from mouse gut was performed with a QIAamp DNA Stool Mini Kit (Qiagen), according to the manufacturer’s instructions. Amplification and sequencing of the 16S rRNA gene was carried out at Genomics unit at Instituto Gulbenkian de Ciência. Samples were amplified using primers specific to the V4 region of the 16S rRNA gene and pair-end sequenced on an Illumina MiSeq Benchtop Sequencer following Illumina recommendations.

### Amplicon sequences processing and analysis

Raw demultiplexed forward and reverse reads were processed using the following methods and pipelines as implemented in QIIME2 version 2018.8 with default parameters unless stated ^21^. DADA2 was used for quality filtering, denoising, pair-end merging and amplicon sequence variant calling (ASV, i.e. sub-OTUs) using qiime dada2 denoise-paired method ^22^. Q20 was used as quality threshold to define read sizes for trimming before merging (parameters: - -p-trunc-len-f and --p-trunc-len-r). Reads were truncated at the position when the 75th percentile Phred score felt below Q20: 235 bp for forward reads and 150 bp for reverse reads. After quality filtering steps, average sample size was 72,908 reads (min: 29,875 reads, max: 103,797 reads). ASVs were aligned using the qiime alignment mafft method ^23^. The alignment was used to create a tree and to calculate phylogenetic relations between ASVs using qiime phylogeny fasttree method ^24^. ASV tables were subsampled without replacement in order to even sample sizes for diversity analysis using qiime diversity core-metrics-phylogenetic pipeline. The smallest sample size was chosen for subsampling. Unweighted and weighted Unifrac distances were calculated to compare community structure ^25^. Taxonomic assignment of ASVs was performed using a Bayesian Classifier trained with Silva V4 database (i.e. 99% OTUs database truncated at position 515 and 826 bp) using the qiime feature-classifier classify-sklearn method ^26^. Differential abundance of taxa was tested using Mann-Whitney (or Kruskal Wallis) non-parametric test on relative abundance of taxa (total sum scale). After Kruskal Wallis, Conover’s test with FDR Benjamini-Hochberg correction is added for pairwise comparison. Unifrac distance matrices and ASV tables were used to calculate principal coordinates and construct ordination plots using R software package version 3.4.3 (http://www.R-project.org). The significance of groups in community structure was tested using Permanova. BiodiversityR version 2.8-4 and vegan version 2.4-5 packages were used. ggplot version 2.2.1 was used for plotting.

## Results

### Gut microbiota composition is distinct in Trem-2 KO mice and is altered during responses to acute hepatotoxic injury

Here we aimed to assess the impact of acute liver injury induced by a single dose of APAP (acetaminophen) on gut microbiota composition. Fecal samples from wild-type and Trem-2 KO mice were collected before (control) and 3 days (D3) after APAP treatment, at the time point of tissue repair (Figure 1A). Standard analysis of 16S rRNA sequencing data using either unweighted or weighted Unifrac phylogenetic distances revealed that Trem-2 KO mice gut microbiota did not clustered with the wild-type (Figure 1B) and indicated that Trem-2 KO mice harbor a distinct microbiota composition profile showing decreased microbiota diversity (Figure 1C). Interestingly, taxonomy analysis at phylum level (Figure 1D) unveiled that untreated Trem-2 KO mice have a significantly increased relative abundance of Verrucomicrobia and Deferribacteres (Figure 1E) while showing a tendency for decreased Tenericutes relative abundance (Figure S1A). The relatively high representation of Verrucomicrobia, a low diversity phylum may partial explains the overall decrease of microbiota diversity in the Trem-2 KO mice.

**Figure 1:**
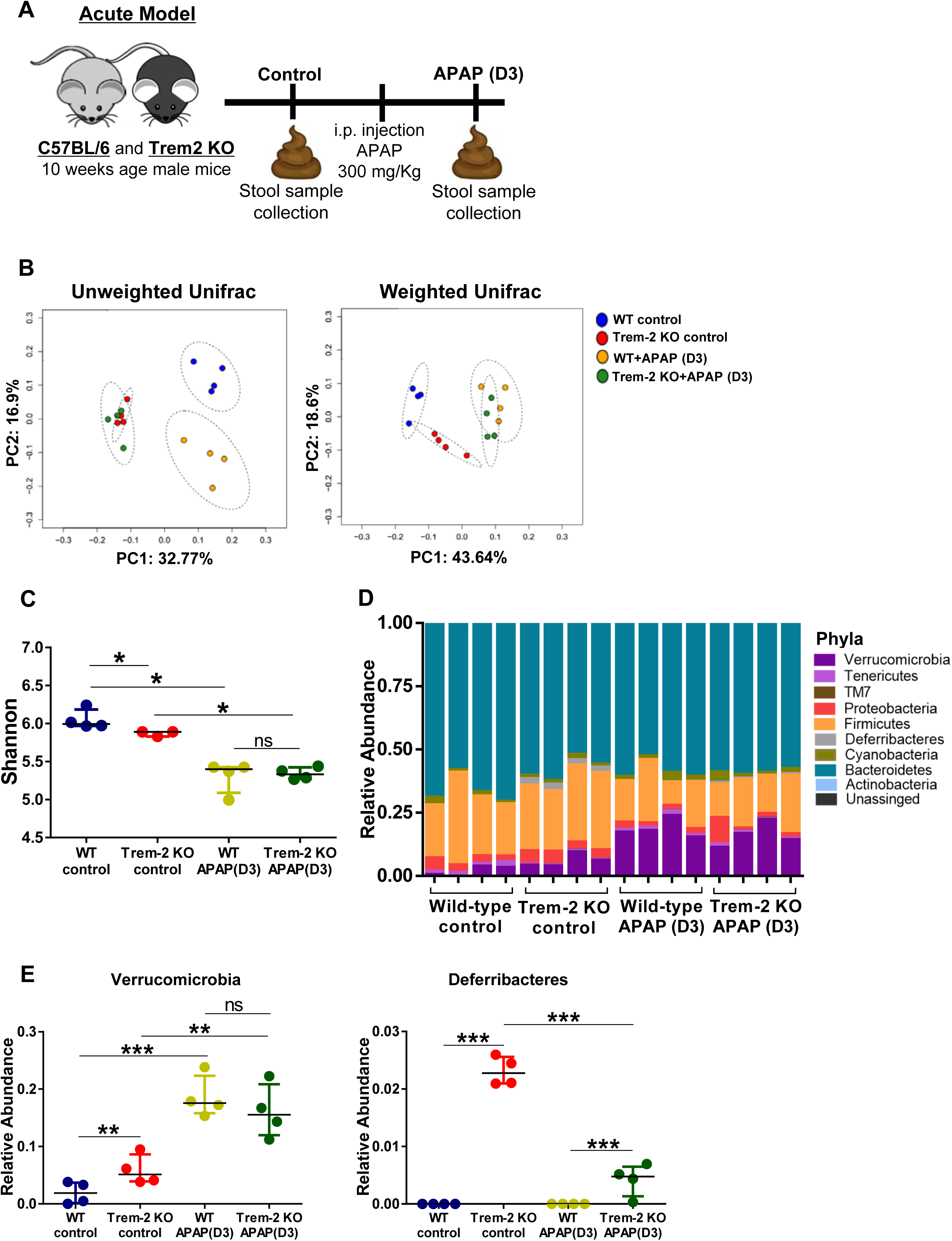
Gut microbiota composition is altered upon acute liver damage induced by APAP. Acute liver damage induced by single intra-peritoneal administration of acetaminophen (APAP) was assessed 3 days (APAP (D3)) post-injection in wild-type and Trem-2 KO mice. Stool samples were collected before (control) and after APAP treatment **(A)**. Principal Coordinate Analysis (PcoA) of 16S rRNA sequencing based on unweighted and weighted Unifrac distances. Each group of mice (WT control, Trem-2 KO control, WT APAP (D3), Trem-2 KO APAP (D3)) is labeled with a different color. Statistics were done using pairwise Permanova test. For unweighted: all groups were statistically different between them, p<0.05. For weighted: WT control vs. Trem-2 KO control and Trem-2 KO control vs. Trem-2 KO APAP (D3) p<0.05 **(B)**. Shannon diversity in the different groups is expressed as the median ± interquartile range. *, p<0.05 using Kruskal-Wallis pairwise analysis **(C)**. Fecal microbiota compositions of each group of mice at phylum level. Each stacked bar represents the microbiota composition of a single mouse for the indicated group. The colored segments represent the relative fraction of each phylum-level taxon **(D)**. Relative abundance of Verrucomicrobia and Deferribacteres phyla for each group, data shown as median ± inter quartile range. *, p<0.05; **, p<0.01; ***, p<0.001 using Conover’s test for multiple comparisons of independent samples **(E)**.

Liver damage induced by APAP treatment was associated with significant changes in microbiota composition occuring both in wild-type and Trem-2 KO mice (Figure 1). The treatment seemed to affect mainly species relative abundance (weighted Unifrac) in Trem-2 KO mice, while wild-type mice are affected both in presence of species (unweighted Unifrac) and their relative abundance (Figure 1B). Overall gut microbiota diversity was decreased in both wild-type and Trem-2 KO when compared to untreated mice (Figure 1C).

Remarkably, APAP treatment induced an increase in Verrucomicrobia, both in wild-type and Trem-2 KO mice (Figure 1D **and** 1E). Although some differences were still identified in phyla with low representation (e.g. Deferribacteres), the APAP treatment had a largely homogenizing effect in the microbiota composition. These results show that untreated Trem-2 KO mice have a distinct microbiota composition profile when compared to the wild-type but most of these differences are lost in mice during acute liver damage induced by APAP treatment.

### Gut microbiota composition is altered during responses to chronic hepatotoxic damage

Given that the drug treatment inducing acute liver damage evoked microbiota alterations, we tested whether a chronic liver insult with a different drug would also affect gut microbiota composition in respect to Trem-2 expression. We collected fecal samples before (control) and after CCl4 treatment, at the time point of fibrosis regression (D3) (Figure 2A). We again observed that untreated Trem-2 KO mice have distinct microbiota composition (Figure 2B, 2D **and** 2E), and decreased microbiota diversity (Figure 2C) as compared to wild-type mice. Remarkably, differences in microbiota composition were recapitulated, with enrichment in Verrucomicrobia and in Deferribacteres phyla in Trem-2 KO mice (Figure 2D **and** 2E) and a significant increase in Tenericutes (Figure S1B).

**Figure 2:**
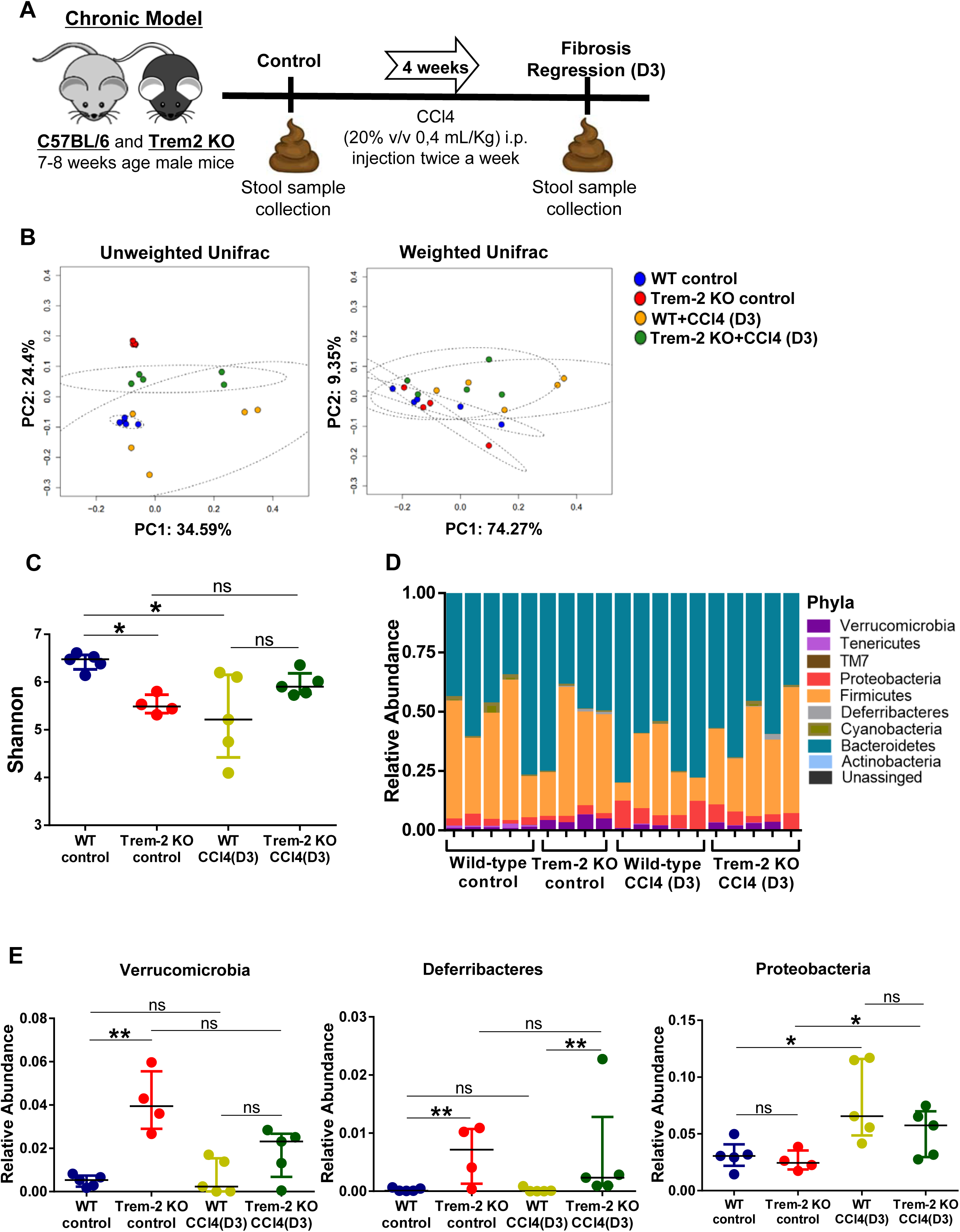
Gut microbiota composition is altered upon chronic liver damage induced by CCl4. Chronic liver injury was induced by administration of CCl4 during 4 weeks, twice a week, in wild-type mice and Trem-2 KO mice. Mice were analyzed three days after the last injection, corresponding to fibrosis regression (D3) time point. Stool samples were collected before (control) and after CCl4 treatment **(A)**. Principal Coordinate Analysis (PcoA) of 16S rRNA sequencing based on unweighted and weighted Unifrac distances. Each group of mice (WT control, Trem-2 KO control, WT CCl4 (D3), Trem-2 KO CCl4 (D3)) is labeled with a different color. Statistics were done using pairwise Permanova test. For unweighted: WT control vs. Trem-2 KO control and Trem-2 KO control vs. Trem-2 KO CCl4 (D3) p<0.05 **(B)**. Shannon diversity in the different groups is expressed as median ± interquartile range. *, p<0.05 using Kruskal-Wallis pairwise analysis **(C)**. Fecal microbiota compositions of each group of mice at phylum level. Each stacked bar represents the microbiota composition of a single mouse for the indicated group. The colored segments represent the relative fraction of each phylum-level taxon **(D)**. Relative abundance of Verrucomicrobia, Deferribacteres and Proteobacteria phyla for each group. Data shown as median ± interquartile range. *, p<0.05; **, p<0.01 using Conover’s test for multiple comparisons of independent samples **(E)**.

Chronic liver injury induced alterations in the microbiota, both in wild-type and Trem-2 KO mice, as shown by principal coordinate analysis of unweighted Unifrac (Figure 2B). No statistical differences were observed between mouse genotypes in weighted Unifrac (Figure 2B) and the chronic treatment blunted differences in microbiota diversity as measured by Shannon diversity index (Figure 2C). Taxonomy analysis showed an increase in relative abundance of Proteobacteria phylum with CCl4 treatment, both in wild-type and Trem-2 KO mice (Figure 2D **and** Figure 2E). These results strongly suggest that independently of the nature of the toxic agent and duration of injury, liver tissue damage is associated with gut microbiota disturbances that decrease species diversity while allowing emergence of specific phyla.

### Gut microbiota composition does not impact on disease outcome in acute liver damage induced by APAP

Finding that Trem-2 modulated the gut microbiota composition in untreated mice prompted us to determine whether this trait was involved in the impaired tissue repair responses observed in Trem-2 KO mice^17^. We performed co-housing experiments followed by administration of APAP. Fecal samples were collected at the beginning of the experiment (control), after wild-type and Trem-2 KO mice have been co-housed for 5 weeks, and at D3 after APAP treatment (Figure 3A). Expectedly, the microbiota composition that was initially different between wild-type and Trem-2 KO, was homogenized by co-housing as shown by unweighted and weighted Unifrac analysis (Figure 3B). Interestingly Trem-2 KO mice tend to acquire a microbiota profile similar to the wild-type. Consistently, Verrucomicrobia phylum predominated in untreated Trem-2 KO mice as compared to the wild-type but this difference was lost after co-housing (Figure 3E **and** 3F). Concerning Tenericutes phylum we observed a tendency for decreased abundance in untreated Trem-2 KO mice that was recovered with co-housing (Figure S2C).

**Figure 3:**
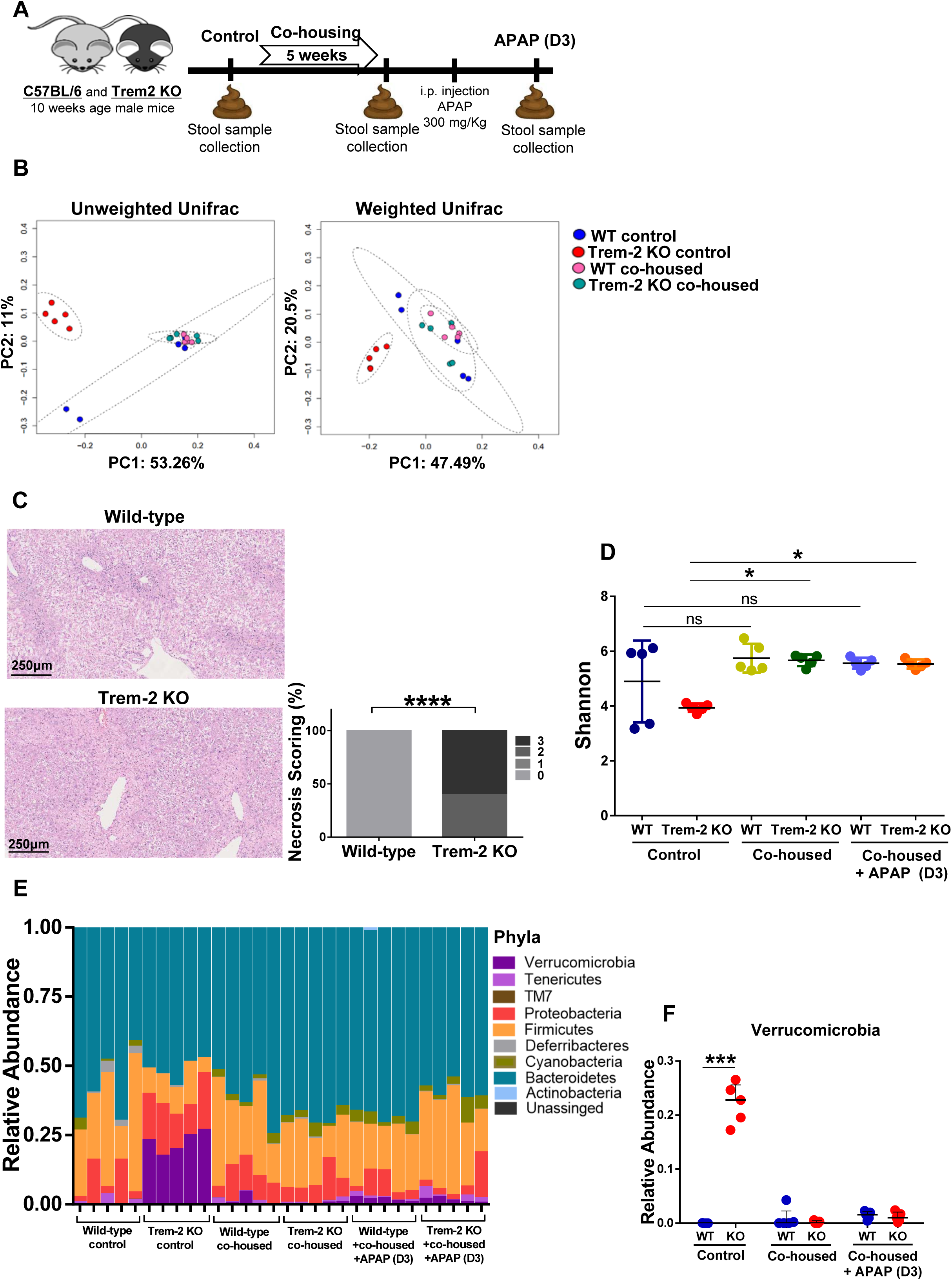
Co-housing experiments homogenize microbiota, but do not impact on acute liver treatment outcome. Wild-type and Trem-2 KO mice were co-housed at 4 weeks of age and were kept together for 5 weeks. After 5 weeks, acute liver damage was induced by single intra-peritoneal administration of acetaminophen (APAP) and disease outcome assessed 3 days (APAP (D3)) post-injection. Stool samples were collected before (control), after co-housing, and after APAP treatment **(A)**. Principal Coordinate Analysis (PcoA) of 16S rRNA sequencing based on unweighted and weighted Unifrac distances. Each group of mice (WT control, Trem-2 KO control, WT co-housed, Trem-2 KO co-housed) is labeled with a different color. Statistics were done using pairwise Permanova test. For unweighted and weighted: WT control vs. Trem-2 KO control and Trem-2 KO control vs. Trem-2 KO APAP(D3) p<0.05 **(B)**. Hepatocyte damage was evaluated by histology in liver sections using hematoxylin & eosin staining and quantified using necrosis scoring. Statistics done using Chi-square test **(C)**. Shannon diversity in the different groups is expressed as median ± interquartile range. *, p<0.05 using Kruskal-Wallis pairwise analysis **(D)**. Fecal microbiota composition of each group of mice at phylum level. Each stacked bar represents the microbiota composition of a single mouse for the indicated group. The colored segments represent the relative fraction of each phyla-level taxon **(E)**. Relative abundance of Verrucomicrobia phyla for each group, shown as median ± interquartile range. ***, p<0.001 using Tukey’s multiple comparisons test **(F)**.

Despite microbiota composition homogenization in the co-housed mice, APAP treatment (D3) recapitulated the hepatic tissue repair phenotype that we have previously described^17^. Namely, wild-type showed complete recover from hepatic tissue damage at D3 after injury, while Trem-2 KO mice revealed persistent liver damage, as quantified by liver necrosis scoring (Figure 3C). These results show that phenotypic differences in wild-type and Trem-2 KO mice during liver regenerative responses after acute injury are not dependent on microbiota composition.

Interestingly, co-housing promoted an increase in microbiota diversity in Trem-2 KO mice, that was similar to wild-type and maintained after APAP treatment (Figure 3D). Intriguingly, microbiota composition was not affected by APAP treatment in co-housed mice (Figure S1A and S1B). This suggests that co-housing may promote alterations in the representation of minor taxa conferring overall microbiota resistance to fluctuations induced by APAP.

Given the role of Verrucomicrobia in mucus production and degradation ^27^ we assessed mucosa fraction and mucus layer in the colon of untreated wild-type and Trem-2 KO mice by histology. We did not find any differences between genotypes (Figure S3), suggesting that over representation of Verrucomicrobia in Trem-2 KO mice is not inducing morphological changes in the mucus layer.

Collectively, these data shows that co-housing leads to microbiota composition homogenization in wild-type and Trem-2 KO mice with increased overall diversity and a drastic decrease of Verrucomicrobia abundance in Trem-2 KO. Additionally, we showed that impaired liver regeneration in Trem-2 KO mice is not dependent on its microbiota composition and that changes induced by co-housing prevail during APAP treatment.

### Trem-2 KO microbiota composition is not restored after co-housing

We next asked whether absence of Trem-2 imposes changes in endogenous gut microbiota composition by analyzing Trem-2 KO and wild-type mice isolated after being co-housed. Fecal samples from wild-type and Trem-2 KO mice were collected before (control) and after 5 weeks of co-housing. The mice were then separated by genotype for an additional period of 5 weeks, after which fecal samples were again collected (Figure 4A).

**Figure 4:**
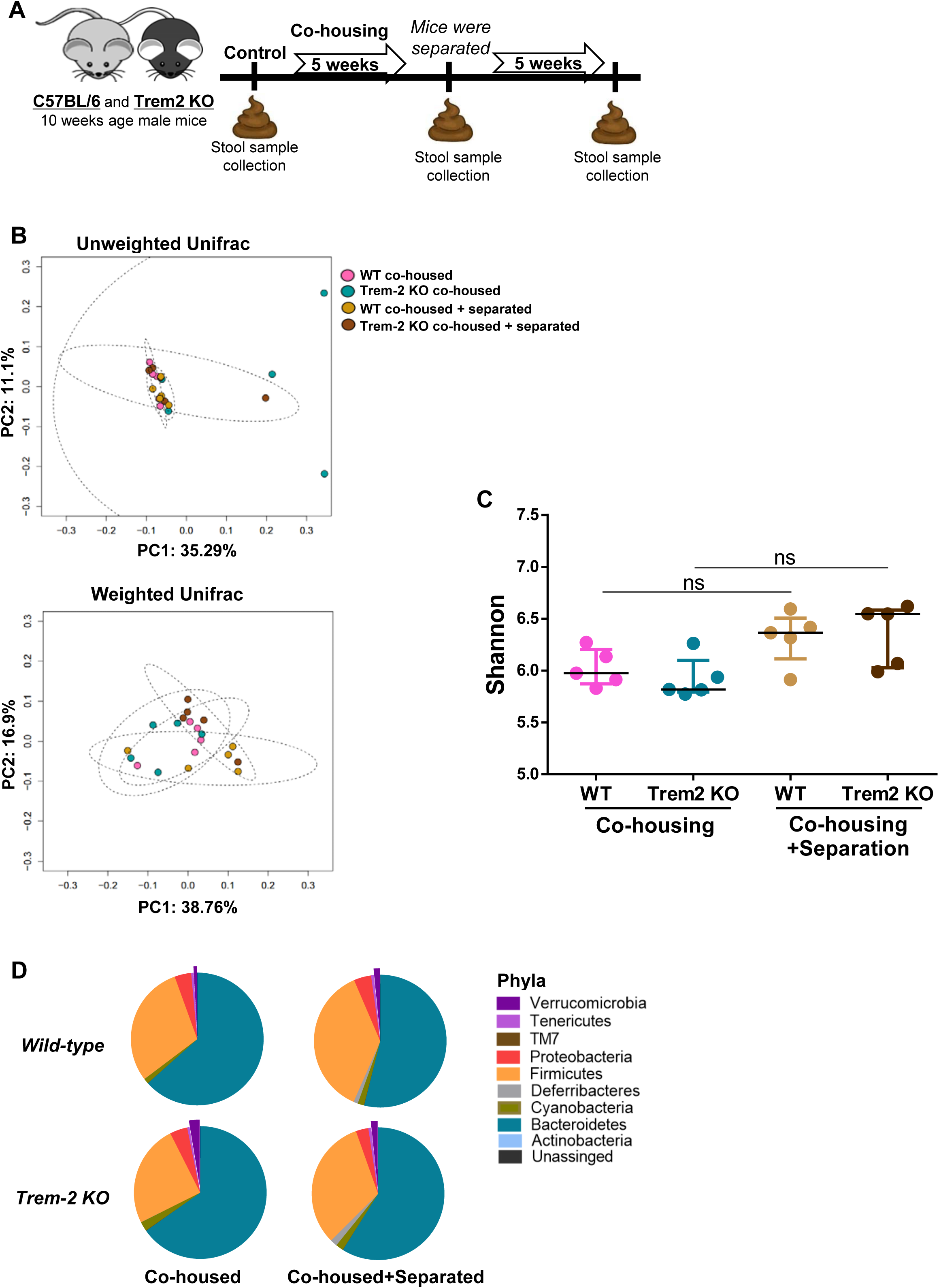
Trem-2 KO does not recover initial microbiota composition after 5 weeks separation post co-housing. Wild-type and Trem-2 KO mice were co-housed at 4 weeks of age and were kept together for 5 weeks. After 5 weeks wild-type and Trem-2 KO mice were separated for 5 more weeks. Stool samples were collected before (control), after co-housing and after mice have been separated **(A)**. Principal Coordinate Analysis (PcoA) of 16S rRNA sequencing based on unweighted and weighted Unifrac distances. Each group of mice (WT co-housed, Trem-2 KO co-housed, WT co-housed+separated, Trem-2 KO co-housed+separated) is labeled with a different color. Statistics were done using pairwise Permanova test **(B)**. Shannon diversity in the different groups is expressed as the median ± interquartile range. Statistics done using Kruskal-Wallis pairwise analysis **(C)**. Relative abundance of the phyla found in the gut microbiota for each genotype after co-housing and after mice have been separated. Each pie chart represents the mean of each experimental group **(D)**.

As shown above (Figure 3) co-housing reproducibly promoted microbiota homogenization between genotypes, measured by unweighted and weighted Unifrac (Figure 4B) and Shannon diversity index (Figure 4C). Remarkably, upon 5 weeks of isolation after co-housing, no differences were observed in microbiota composition and diversity when comparing wild-type and Trem-2 KO mice (Figure 4B **and** 4C). Moreover, phyla composition was maintained (Figure 4D) and Verrucomicrobia did not change after separation.

These results suggest that co-housed Trem-2 KO mice do not have the ability to restore their initial microbiota even after 5 weeks of separation. Thus Verrucomicrobia phylum seems to be easily outcompeted in the presence of other bacteria species that were probably absent in Trem-2 KO mice before co-housing.

## Discussion

This work evaluated gut microbiota composition in two different experimental models of liver regeneration, induced acutely by APAP or chronically by CCl4 administration in wild-type and Trem-2 KO mice. We showed that both treatments lead to gut microbiota changes, with an overall decrease in diversity. Specifically, we observed an enrichment of Verrucomicrobia phylum in the acute model and an increase in Proteobacteria phylum in the chronic model, irrespective of Trem-2 genotype. Interestingly, despite significant differences in microbiota composition between untreated wild-type and Trem-2 KO mice, liver damage indistinctly affected the gut microbiota leading to similar diversity in both mouse genotypes.

Previous reports have shown that microbiota is changed in liver disease^28^ and associated pathologies such as non-alcoholic fatty liver disease (NAFLD), cirrhosis^28^, type 2 diabetes and obesity^29, 30^. Different studies reported associations between Verrucomicrobia and liver related diseases. Verrucomicrobia comprises the *Akkermansia* genus which is a mucus degrading bacteria using mucins as energy source, highly rich in aminoacids and oligosaccharides^27^.

In obese subjects abundance of *Akkermansia muciniphila* is associated with a healthier metabolic status and better clinical outcomes after caloric restriction^31^. The same association was observed in mice fed with high fat diet (HFD)^32^ and in patients with type 2 diabetes^32, 33^. Moreover it was shown that administration of a purified membrane protein form *Akkermansia muciniphila* improves metabolism in obese and diabetic mice^34^. Interestingly, usage of metformin, an antidiabetic drug, has been shown to increase levels of *Akkermansia* spp.^35^. Overall, there is a strong association between increased representation of Verrucomicrobia and improved metabolic control in liver disorders. In addition, it is consensual that in obesity and type 2 diabetes there is lower gut bacterial diversity^30^.

Surprisingly we found that untreated Trem-2 KO mice have significant reduction in microbiota diversity with a striking increase in Verrucomicrobia abundance. Trem-2 involvement in Alzheimer’s disease^36^ and multiple sclerosis^16^ is well explored. Incidentally, different studies show an over representation of Verrucomicrobia phylum in neurodegenerative diseases^37, 38^. This body of evidence raised the possibility that the modulation of gut microbiota by Trem-2 here reported, may be involved in the association of Trem-2 with metabolic and neurodegenerative disorders.

Co-housing experiments revealed that Trem-2 KO mice homogenize their microbiota to a similar composition as found in wild-type, suggesting that wild-type microbiota has competitive advantage. Remarkably, the initial Verrucomicrobia abundance found in Trem-2 KO mice is not recovered upon long-term separation after co-housing. This suggests that Verrucomicrobia in untreated Trem-2 KO mice have indeed reduced competitive advantage when in the presence of a richer microbiota community. A study where two subjects showed an 80% increase of *Akkermansia* spp. after treatment with broad-spectrum antibiotics ^39^, supports that Verrucomicrobia has preferential expansion in low competitive gut microbiota environment.

Furthermore, we unveiled that the microbiota composition in Trem-2 KO mice is skewed to a profile that resembles cases of dysbiosis with over representation of Verrucomicrobia. Nevertheless, we show that such microbiota profile is not a determining factor underlying tissue repair impairments in the Trem-2 KO. A study using antibiotics administration followed by CCl4-induced acute liver injury showed that gut microbiota removal has a protective effect on Trem-2 KO mice, as shown by decreased hepatic enzymes in the serum of these mice^18^. Moreover in a mouse model of renal ischemia-reperfusion injury, depletion of gut microbiota using broad-spectrum antibiotics significantly attenuated renal damage and dysfunction, which was associated to lower levels of F4/80 expression and of CX3CR1 and CCR2 chemokines in resident macrophages and in bone-marrow monocytes^40^. In our study we did not observe an impact of gut microbiota in liver injury outcomes under microbiota perturbations imposed by Trem-2 KO. Taken together these studies suggest that responses to liver damage induced by hepatotoxic drugs are affected by depletion of gut microbiota but we demonstrate that the mere gut microbiota perturbations with increased Verrucomicrobia representation is not sufficient to determine impairments in response to liver damage observed in Trem-2 KO mice.

Nevertheless, our results corroborate previous reports showing that liver damage induces gut microbiota composition alterations, mostly by imposing decreased diversity. Further investigations are needed to determine if microbiota alterations associated with liver damage play a subsequent pathogenic role in the development of other organic and systemic disorders, particularly when associated with emergence of Verrucomicrobia *Akkermansia* spp.

## Acknowledgements

The authors acknowledge the histology, genomics and bioinformatics units at IGC, in particular Dr. Belen Carbonetto for assistance with data analysis.

---

**Supplementary Figure 1:**
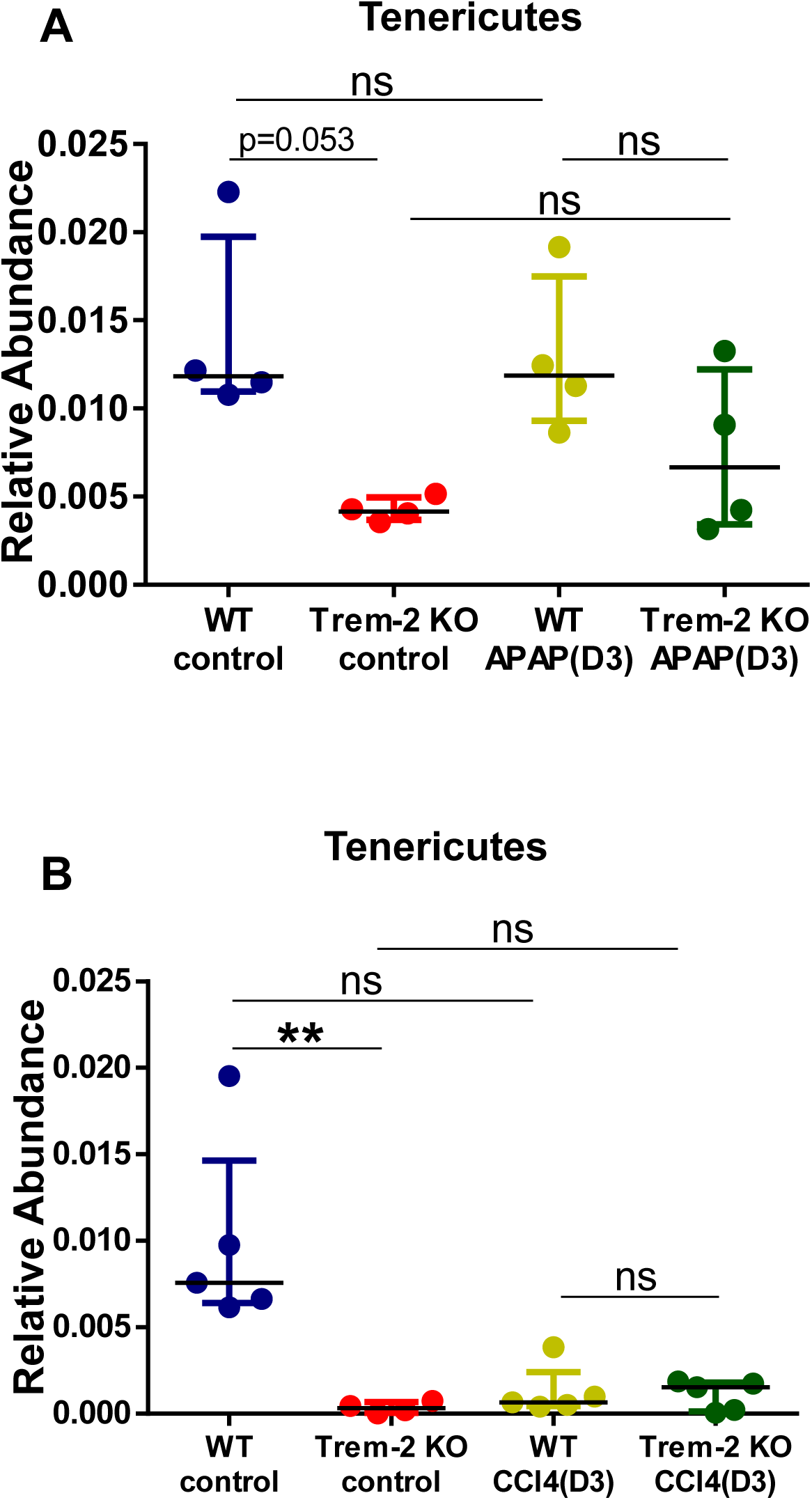
Relative abundance of Tenericutes in acute and chronic liver models. Relative abundance of Tenericutes phyla for each group in wild-type and Trem-2 KO mice shown as median ± interquartile range in the acute **(A)** and in the chronic **(B)** models. **, p<0.01 using Tukey’s multiple comparisons test.

**Supplementary Figure 2:**
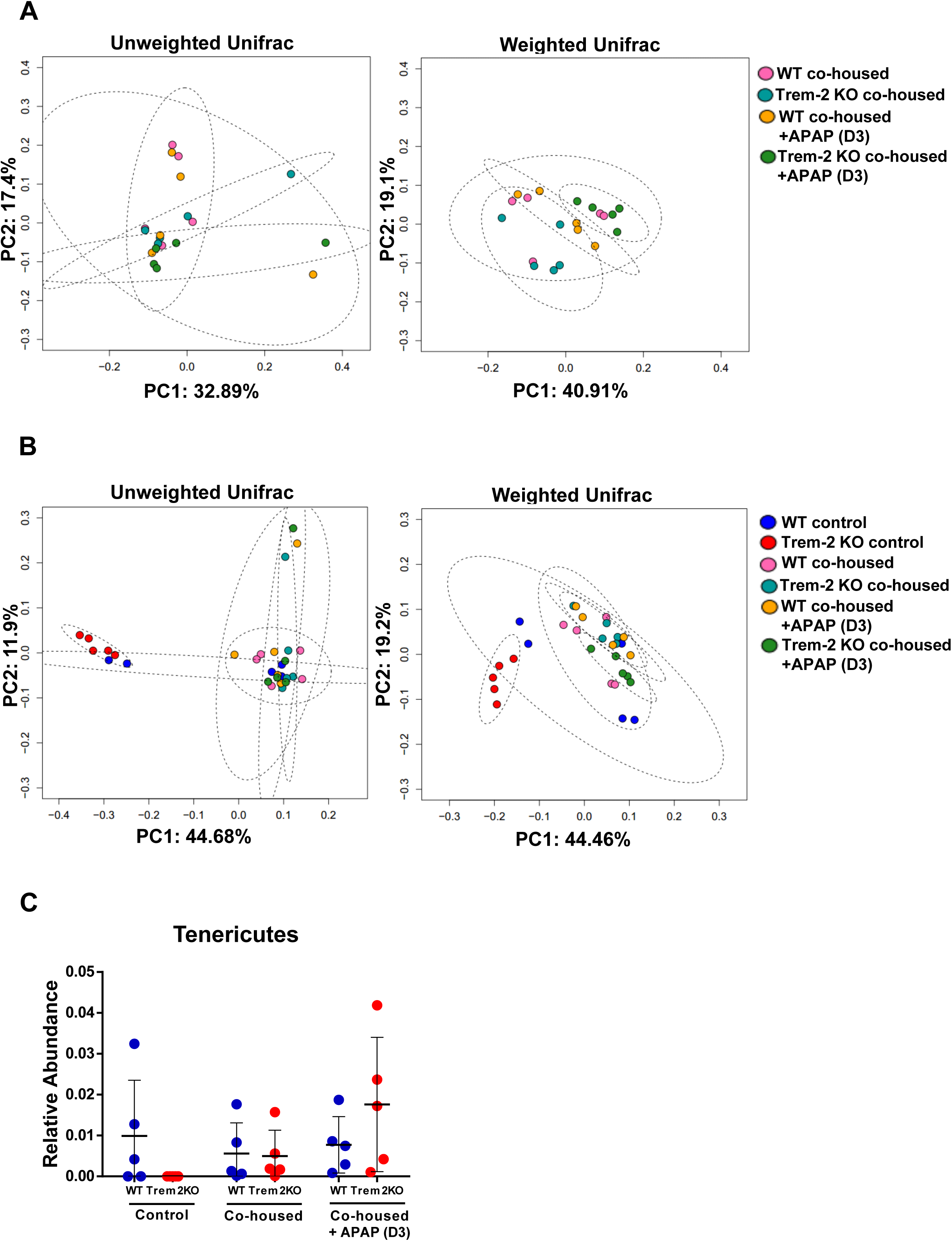
Co-housing experiments promote microbiota homogenization between wild-type and Trem-2 KO mice. Principal Coordinate Analysis (PcoA) of 16S rRNA sequencing based on unweighted and weighted Unifrac distances for mice that were co-housed and consequently treated with APAP. Each group of mice (WT co-housed, Trem-2 KO co-housed, WT co-housed+APAP, Trem-2 KO co-housed+APAP) is labeled with a different color **(A)**. Principal Coordinate Analysis (PcoA) of 16S rRNA sequencing based on unweighted and weighted Unifrac distances for all the groups represented together. Each group of mice (WT control, Trem-2 KO control, WT co-housed, Trem-2 KO co-housed, WT co-housed+APAP, Trem-2 KO co-housed+APAP) is labeled with a different color **(B)**. Relative abundance of Tenericutes phyla for each group shown as median ± interquartile range **(C)**. Statistics used pairwise Permanova test in A and B and using Tukey’s multiple comparisons test C.

**Supplementary Figure 3:**
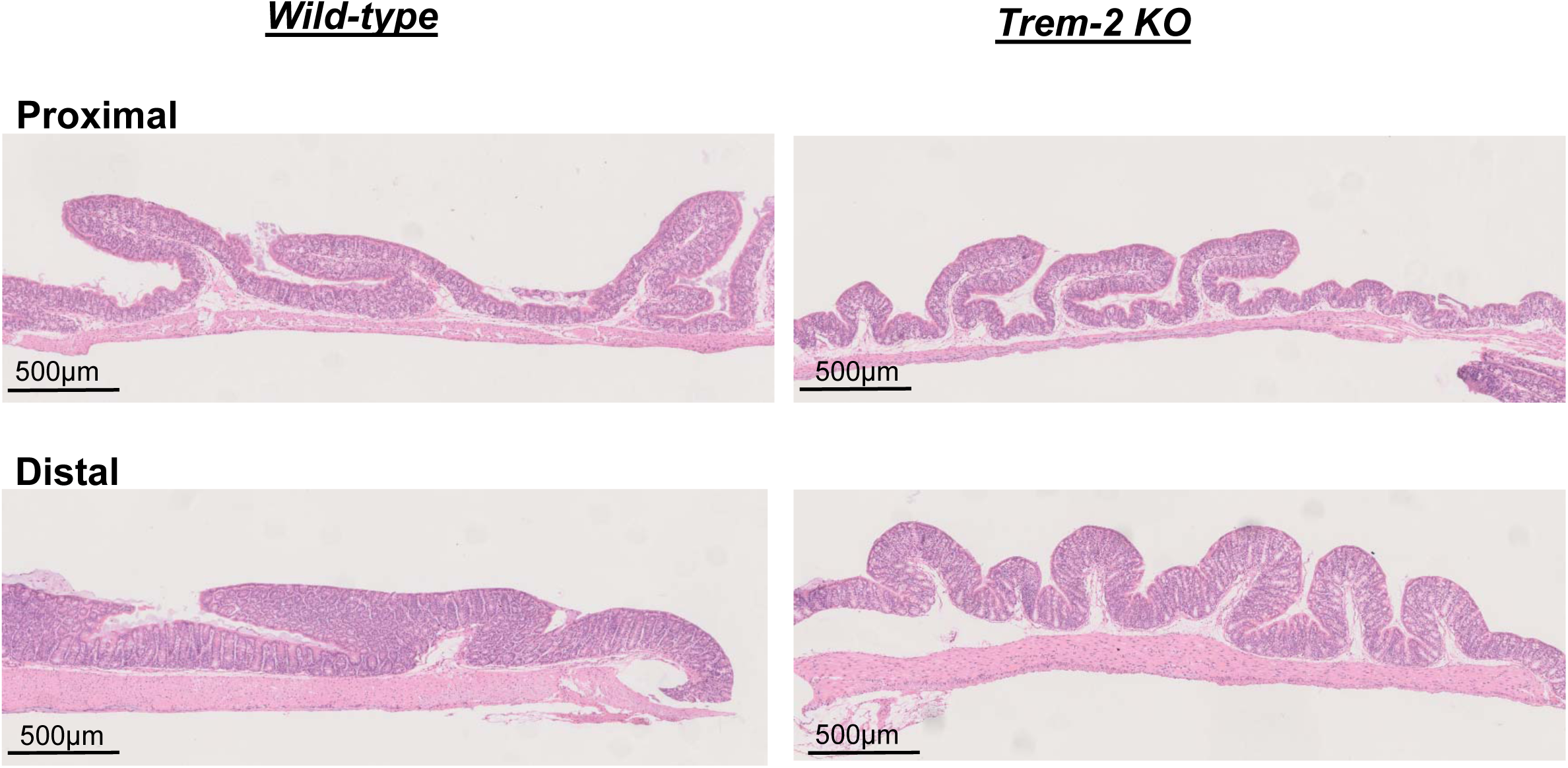
Histological analysis of intestine from untreated wild-type and Trem-2 KO mice. Hematoxylin-eosin staining of paraffin-embedded sections of the colon. Proximal and distal colon parts are shown for untreated wild-type and Trem-2 KO mice.

